# Impulse perturbation reveals cross-modal access to sensory working memory through learned associations

**DOI:** 10.1101/2023.03.01.530587

**Authors:** Güven Kandemir, Elkan G. Akyürek

**Author notes:** Address correspondence to: Güven Kandemir, Institute for Brain and Behavior, Vrije Universiteit Amsterdam Van der Boechorststraat 7, 1081 BT Amsterdam, The Netherlands.

## Abstract

We investigated if learned associations between visual and auditory stimuli can afford full cross-modal access to working memory. Previous research using the impulse perturbation technique has shown that cross-modal access to working memory is one-sided; visual impulses reveal both auditory and visual memoranda, but auditory impulses do not seem to reveal visual memoranda (Wolff et al., 2020b). Our participants first learned to associate six auditory pure tones with six visual orientation gratings. Next, a delayed match-to-sample task for the orientations was completed, while EEG was recorded. Orientation memories were recalled either via their learned auditory counterpart, or were visually presented. We then decoded the orientation memories from the EEG responses to both auditory and visual impulses presented during the memory delay. Working memory content could always be decoded from visual impulses. Importantly, through recall of the learned associations, the auditory impulse also evoked a decodable response from the visual WM network, providing evidence for full cross-modal access. We also observed that after a brief initial dynamic period, the representational codes of the memory items generalized across time, as well as between perceptual maintenance and long-term recall conditions. Our results thus demonstrate that accessing learned associations in long-term memory provides a cross-modal pathway to working memory that seems to be based on a common coding scheme.

## Introduction

Working memory (WM) is the ability of an organism to retain and manipulate information over a brief period of time in the absence of sensory input (Baddeley, 2000). The neural basis of WM has recently been a topic of great interest. For instance, accumulating evidence shows that WM uses the sensory cortices to retain low-level sensory stimuli (Harrison & Tong, 2009; Kumar et al., 2016; Serences et al., 2009, Rademaker et al., 2019; but see also Bettencourt & Xu, 2016). It has also been discovered that working memory maintenance can take place without ongoing observable neural activity (Lewis-Peacock, et al., 2012; LaRocque et al., 2013). Such activity-silent or quiescent maintenance of information may be supported by transient alterations in the synapses of the memory network (i.e., short-term synaptic plasticity; Mongillo et al., 2008). Connecting these findings is recent evidence from studies using a visual impulse perturbation technique that have provided some support for the notion that such quiescent storage may occur in occipital regions (Wolff et al., 2015; 2017; 2020a).

The visual impulse perturbation technique entails the presentation of a salient, but invariant and task-irrelevant stimulus during WM maintenance. The brain’s response to this impulse stimulus should also be invariant, apart from possible interactions caused by the presence of memoranda in the WM network, such as might be instantiated by changes in synaptic connectivity (Stokes, 2015). In a series of studies by Wolff and colleagues (2015; 2017; 2020a), this proved to be the case, and visual WM contents could be decoded from the impulse response, as measured by EEG over posterior regions.

Although the impulse perturbation technique is primarily a neuroscientific tool for scientists to read out what is being remembered in the brain, it may also be an analogue of how the brain itself performs memory read-out. In order to retrieve a memory, all that might be needed is a generic, self-evoked elevation of activity in the relevant network, and the ‘impulse response’ would produce the memoranda. In this context, it is important to consider the extent of the network. Are WM contents modality-specific, locked in their respective sensory cortices? Or is there a modality-unspecific pathway, paralleling the cross-modal neural interactions that have been reported between audition and vision (Martuzzi et al., 2007; Marian et. al., 2021; Thelen & Murray, 2013, Thelen et al., 2014; 2015)?

Evidence from Wolff and colleagues (2020b) suggests that the memory network can indeed be perturbed cross-modally, but not equally so for the auditory and the visual working memory network. In their study, Wolff and colleagues asked participants to memorize auditory pure tones or visual orientation gratings in different blocks. A retro-cue indicated the task-relevant item. During the delay period following the cue, two task-irrelevant impulse signals were presented; one auditory and one visual. From both visual and auditory impulse-driven EEG activity the auditory stimuli could be successfully decoded. However, this bi-modal impact of the visual impulse was not mirrored by the auditory impulse, which revealed auditory WM contents, but not visual ones.

This difference in the impulse response between the modalities could stem from the general dominance of the visual system (Martuzzi et al., 2007; Posner et al., 1976). Alternatively, it might be due to differences in the strength or utility of the connections between the auditory and the visual sensory cortices (Eckert et al., 2008). In the latter case, it might be hypothesized that these connections might be plastic and could be modulated, for instance through learning, which might open up a new access pathway to WM. Support for this hypothesis comes from animal studies, which have shown that via classical conditioning, auditory stimuli can alter the activation level of the visual cortex (Ibrahim et al., 2016; Garner & Keller, 2021; Petro et al., 2017). The current study was set up to investigate this possibility.

In our experiment, we created direct links between auditory and visual items by having our participants learn to associate pairs of tone and orientation stimuli. We hypothesized that these learned, long-term associations would enable a new cross-modal pathway to WM from the auditory to the visual modality. To preview the outcomes, we replicated and extended the visual impulse effect, showing that the response to a visual impulse revealed visual memory contents, both when this visual content was actually presented during the trial, and when it was retrieved from long-term memory through its auditory pair. The associated EEG patterns for the orientations cross-generalized, suggesting a common coding scheme. Importantly, we found that the auditory impulse also revealed visual memories, but only when these visual memories were retrieved from long-term memory via their auditory pairs. The results suggest that access paths to WM may be plastic and amenable to alterations through learning.

## Method

### Participants

Thirty healthy adults (22 female, M_Age_ = 24.67, s.d._Age_ = 5.01) volunteered in the EEG sessions in return for monetary compensation. This final sample of participants was selected from a larger group on the basis of a pre-screening test, which assessed memory retrieval of the learned auditory-visual stimulus pairs by using a shorter version of the experimental task with 288 trials (recall condition, see below). The selection criteria were 70 % overall accuracy and average response speed below a second. Participants were fully informed about the aim and the design of the experiment and provided written consent for participation and the usage of the data. The study was approved by the Ethical committee of the University of Groningen (ID Code = PSY-1920-S-0092).

### Stimuli and Apparatus

In the EEG experiment, stimulus presentation was controlled by the freely available Matlab extension Psychtoolbox (Brainard, 1997; Kleiner et al., 2007). Throughout the experiment a gray background (RGB = 128, 128, 128) was maintained, and a fixation dot consisting of a white circle and a black dot (0.2° visual angle) was always present at the center of the screen. A custom two-button response box with USB connection was used for response collection.

The auditory stimuli that were paired with the visual memory items were 6 pure tones with frequencies ranging from 270 Hz to 1527.4 Hz with half an octave difference in between. All tones contained a 10 ms ramp-up at the beginning and a 10 ms ramp-down at the end. The auditory impulse was created by overlapping 8 pure tones with frequencies ranging from 270 Hz to 3055 Hz with half an octave difference in-between, as was previously done in an earlier study (Wolff et al 2020b). All auditory stimuli were generated with the freely available software Audacity (Audacity Team, 2016). The auditory stimuli were delivered via Sweex in-ear headphones. The sound intensity of all auditory stimuli was set to 75 dBs on average with a deviation of 2 dB, according to a Tenma 72-860 sound level meter measured from the right side of the headphones. A Focusrite external soundcard was used for the presentation of the auditory stimuli to ensure accurate timing.

The visual memory items consisted of 6 discrete sine-wave gratings with orientations ranging from 15° to 165°, spaced in steps of 30°. The probes were sine-wave gratings and always deviated from the memory items by (−/+) 4°, 9°, 15°, 22°, 30° or 39°, in clockwise or counter-clockwise direction. Probes were uniformly distributed. All gratings were presented with 0.5 contrast and had a diameter of 6.46°. The spatial frequency of the gratings was 1 cycle per degree and the phase of the gratings was randomized in each trial. The visual impulse consisted of three white disks, with a diameter of 9.69°. They were centered on the horizontal axis, and overlapped by half their diameter. All visual stimuli were presented at the center of a 17 inch (43.18 cm) Samsung CRT monitor with a resolution of 1024 by 768 pixels and 100 Hz refresh rate.

### Procedure

Prior to the EEG experiment, participants learned auditory-visual stimulus pairs in a location of their own choosing, on their personal devices, and at their own pace by using the freely available OpenSesame program (Mathôt et al., 2012). Further details of the self-study procedure are given in the Supplementary Materials.

In the EEG experiment participants completed 2304 trials, divided into 96 blocks, and spread over 4 consecutive sessions, all of which were completed on the same day. Trial flow within a block was automatized, whereas participants could rest between blocks. The main breaks were between sessions, and the duration of these breaks was determined by the participants themselves to counter-balance fatigue effects. The entire experiment lasted approximately 5,5 hours, including the EEG cap preparation and the breaks.

The first two sessions were always the “Recall” sessions, during which an auditory stimulus (i.e., one of the pure tones) was presented to cue for the recall of the memorized orientations (1152 trials), whereas the final two sessions were always “Maintain” sessions, in which the orientation grating was directly presented instead. Session order was constant to avoid extinction of the learned associations between auditory and visual stimuli. For each participant the auditory and visual stimuli were randomly paired without repetition, but with the limiting rule that each auditory stimulus was paired five times with the same visual stimulus over the entire participant sample. This was both to prevent decoding to pick up on a subset of orientations that might be paired with easier-to-remember sounds (e.g., at the low or high end of the frequency range), and to dissociate the circular relationship between orientation stimuli from the ordinal frequency differences between auditory cues.

Participants were seated in a well-lit room approximately 60 cm away from the monitor and were instructed to keep their gaze fixated on the fixation dot that was presented at the center of the screen at all times. The first trial of each block was initiated by pressing the Spacebar, after which a “Get Ready” message was presented in 12 point Arial font for 1000 ms. Next, the fixation dot was presented for 700 ms until the onset of the memory item, which was displayed for 200 ms (Fig. 1). In the first two sessions this item was one of the six auditory tones, whereas in the final two sessions it was one of the visual orientation gratings.

**Figure 1.**
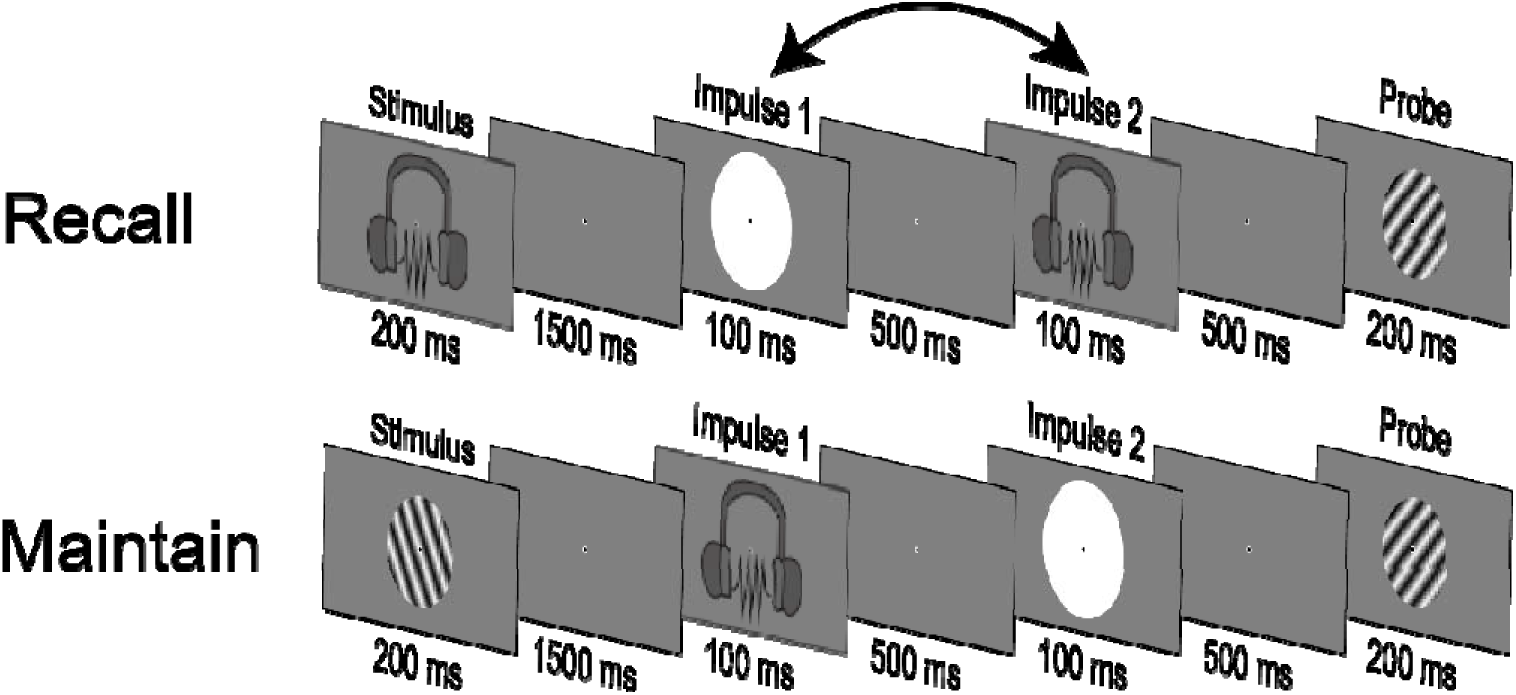
Experimental trials in the Recall (top) and Maintain (bottom) conditions from memory item onset to response probe.

The memory item was followed by a delay of 1500 ms, during which participants were expected to recall the orientation that was paired with the auditory tone or maintain the orientation of the visually presented orientation grating. In order to ensure that participants kept their gaze at the fixation dot prior to impulse, 1000 ms after memory item onset the fixation dot disappeared for 100 ms and re-appeared again. At the end of the post-memory item delay, the first impulse signal (auditory or visual) was presented for 100 ms, followed by a delay with 500 ms duration. Afterwards, the second impulse (which was always in the other modality than the first impulse) was presented for 100 ms, again followed by a delay of 500 ms duration. The order of the impulse modalities was maintained throughout each session, but was swapped between sessions, and the order was counterbalanced within and across participants.

Next, the probe was presented for 200 ms, followed by a blank screen with a fixation dot that was shown until a response was given. Participants had to press the left or the right button on the response box to indicate the direction of the deviation of the probe relative to the memory orientation. After the response, a smiley was presented for 300 ms to provide accuracy feedback. At the end of each block participants were presented with a graph showing their accuracy in that block, and overall, as well as a graph showing their response time. Participants were always encouraged to respond both fast and accurately.

### EEG Acquisition and Preprocessing

The EEG was recorded with Brainvision recorder software and an analog-to-digital TMSI Refa 8-64/72 amplifier, from 62 Ag/AgCl sintered electrodes deployed according to the extended international 10-20 system. The data was acquired at a 1000 Hz sampling rate, and referenced online to the average of all electrodes. An electrode above the sternum served as ground. Vertical EOG was recorded bipolarly from electrodes placed superior and inferior to the left eye. Horizontal EOG was recorded from electrodes placed lateral to each eye. The impedance at all electrodes was retained below 5Ωks throughout the experiment.

The data was downsampled to 500 Hz and filtered offline using a band-pass filter (0.1 Hz high pass – 40 Hz low pass) using the freely available Matlab extension EEGlab (Delorme & Makeig, 2004). Next, the data was epoched relative to the onset of the memory items (−200 ms – 1500 ms) and relative to Impulse 1 and 2 (−200 – 600 ms). The epochs were investigated for artefacts by visualizing the data with the freely available Matlab extension Fieldtrip (Oostenveld et al., 2010). Epochs with blinks, muscle artefacts and extensive noise were selected and removed from the analyses. Noisy channels were replaced by spherical spline interpolation. The average percentage of removed trials for each epoch was as follows: 17.88 % in the memory item epoch (Recall and Maintain condition together), 10 % in the Impulse 1 epoch (auditory and visual together), and 9.18 % in the Impulse 2 epoch (auditory and visual together).

### Multivariate Analyses

All analyses were conducted over 17 posterior electrodes (‘P7’, ‘P5’, ‘P3’, ‘P1’, ‘Pz’, ‘P2’, ‘P4’, ‘P6’, ‘P8’, ‘PO7’, ‘PO3’, ‘POz’, ‘PO4’, ‘PO8’, ‘O2’, ‘O1’, ‘Oz’), unless explicitly stated otherwise. In time-course decoding, the EEG data was first baselined to the average of a −200 to 0 ms time window, relative to stimulus onset. Next the data was further down-sampled to 125 Hz to reduce computation time. The data were entered into a multivariate decoding analysis with 8 fold cross-validation. The training set consisted of 7 folds of the data. First, trials in the training set were distributed over 6 bins, based on their orientations. In order to avoid any biases, the trial count in each bin was equalized by randomly subsampling trials in each bin as many times as the number of trials in the smallest bin. The EEG data in each bin was averaged over trials, yielding a multi-variate pattern for each condition. Next, each trial in the test set was contrasted with the averaged patterns at each time point, providing a measure of similarity. Similarity was quantified by means of Mahalanobis distance (De Maesschalck et al., 2000), and the covariance matrix was estimated from the entire training set using a shrinkage estimator (Ledoit & Wolf, 2004). In order to ease interpretation, the sign of the distance values were reversed, so that higher similarity meant larger values. The distance measures were then mean-centered and convolved with the cosine function of the memory orientations, providing a single decoding accuracy output at each time point in each trial. This procedure was repeated 8 times so that all folds served as test set. The entire decoding procedure was repeated 100 times in order to avoid spurious variations caused by random sampling of the training set. The resulting decoding output was averaged over repetitions. The results for each trial were averaged and smoothened with a Gaussian filter (kernel =2) for each participant at each time point.

The temporal generalization of the neural code was assessed by training on each specific time point within an epoch and testing this on all possible time points. This was only applied to conditions and epochs with statistically significant decoding in order to assess the temporal stability of the representations. Evidence for dynamic coding was defined as higher decoding accuracy when training and testing at the same time point, relative to testing on other time points. The preparation of the EEG data and the decoding procedure were identical to time-course decoding.

To assess the generalization of representations from one condition to another, the training and the testing data were selected from separate conditions. In order to acquire a single decoding accuracy measure per trial, the EEG data was pooled over space and time dimensions as in earlier studies (e.g., Wolff et al., 2020a), to focus the analyses on the fast dynamics of the ERPs. The critical period was defined as the 300 ms window ranging from 100 to 400 ms relative to the impulse signal, as in earlier studies (Wolff et al 2017; 2020a, 2020b). First, the mean activity within the critical time window was subtracted from the entire window in order to remove stable EEG activity. This served as a baseline as well. Next the EEG voltages on each electrode were down-sampled by using a 10 ms moving average. The remaining voltages were expanded over electrode space as features, such that each trial contained a single spatio-temporal pattern consisting of 510 values (30 time-points over 17 electrodes). Similar to time-course decoding, this data was decoded using 8 fold cross-validation, and Mahalanobis distance was the measure of similarity. Critical time-period decoding was also repeated 100 times to account for variations due to sampling and the final results were averaged over all repetitions and trials, yielding a single decoding accuracy for each participant.

Visualization of memory representations in state space was accomplished by multi-dimensional scaling of the pairwise differences between each item in each condition. For this, data was baselined and pooled over time and electrode space using the 100-400 ms window in the same way as in the critical-window approach explained above (Wolff et al., 2020a). Next, an equal number of trials was subsampled for each memory item in each condition. The pooled data in each bin was averaged, forming item-specific patterns in each condition. Next, pairwise Mahalanobis distances were calculated, by using the correlation-covariance matrix estimated using the entire data set. This procedure was repeated 100 times for each participant. The distance values were averaged over all repetitions and participants. The averaged values were then used to estimate the relative positions of representations in three-dimensional state-space.

The topographical distribution of the signal associated with the memory items was analyzed with an EEG application of the searchlight technique (van Ede et al, 2019; Wolff et al., 2020a; 2020b). Data preparation was identical to the critical-window analysis explained above, with the exclusion that instead of using 17 electrodes to pool over, each electrode and its closest two neighbors were used in each analysis to predict the trial-specific orientation. This procedure was repeated 100 times for all 62 electrodes. By averaging over repetitions and trials a single decoding value was created for each electrode for each participant, and the average decoding accuracy over all participants was visualized using Fieldtrip (Oostenveld et al., 2010).

### Statistical analysis

Decoding accuracy values of all participants were analyzed with non-parametric sign permutation tests. The sign of the mean decoding accuracy that was averaged over trials for each participant was randomly flipped 100 000 times with 0.5 probability and the formed group means were used to build a null distribution. The proportion of the mean decoding accuracy relative to this distribution was reported as the *p* value, for which the cut-off for statistical significance was 0.05. For time-course data, the same statistical assessment was done at each time-point with cluster correction. For the topographical output, the decoding accuracy of each electrode was assessed with the same group permutation test and the electrodes that contributed significantly (*p* < 0.05) were presented.

The statistical assessment of the temporal generalization was based on the difference of the decoding accuracy on the diagonal (train and test on the same exact time point) from the horizontal axis (tested on all possible time points). This was tested against zero with a permutation test (n= 100 000) with a cluster correction. Values above the cut-off threshold (*p* < 0.05) were considered indicative of a dynamic neural code, as they reflect significantly better decoding on the diagonal axis. Cross-generalization between conditions was assessed by a sign permutation test to decide if the decoding accuracy was higher than zero, similar to the time course analyses.

## Results

### Behavioral Results

Accuracy was worse when orientations were retrieved from long-term memory in Recall trials, relative to Maintain trials (Fig. 2A; *t*(29) = 8.878; *p* < 0.001, paired; *M _Maintain_* = 0.822, *s.d. _Maintain_* = 0.061, *M _Recall_* = 0.753, *s.d. _Recall_* = 0.05). Response accuracy varied predictably as a function of probe deviation, but performance was worse in Recall trials than in Maintain trials for all probe-item deviations (Fig. 2B, non-parametrical permutation test *n _Perm_* = 100 000, *p* < 0.001, *two-tailed, cluster corrected*).

**Figure 2.**
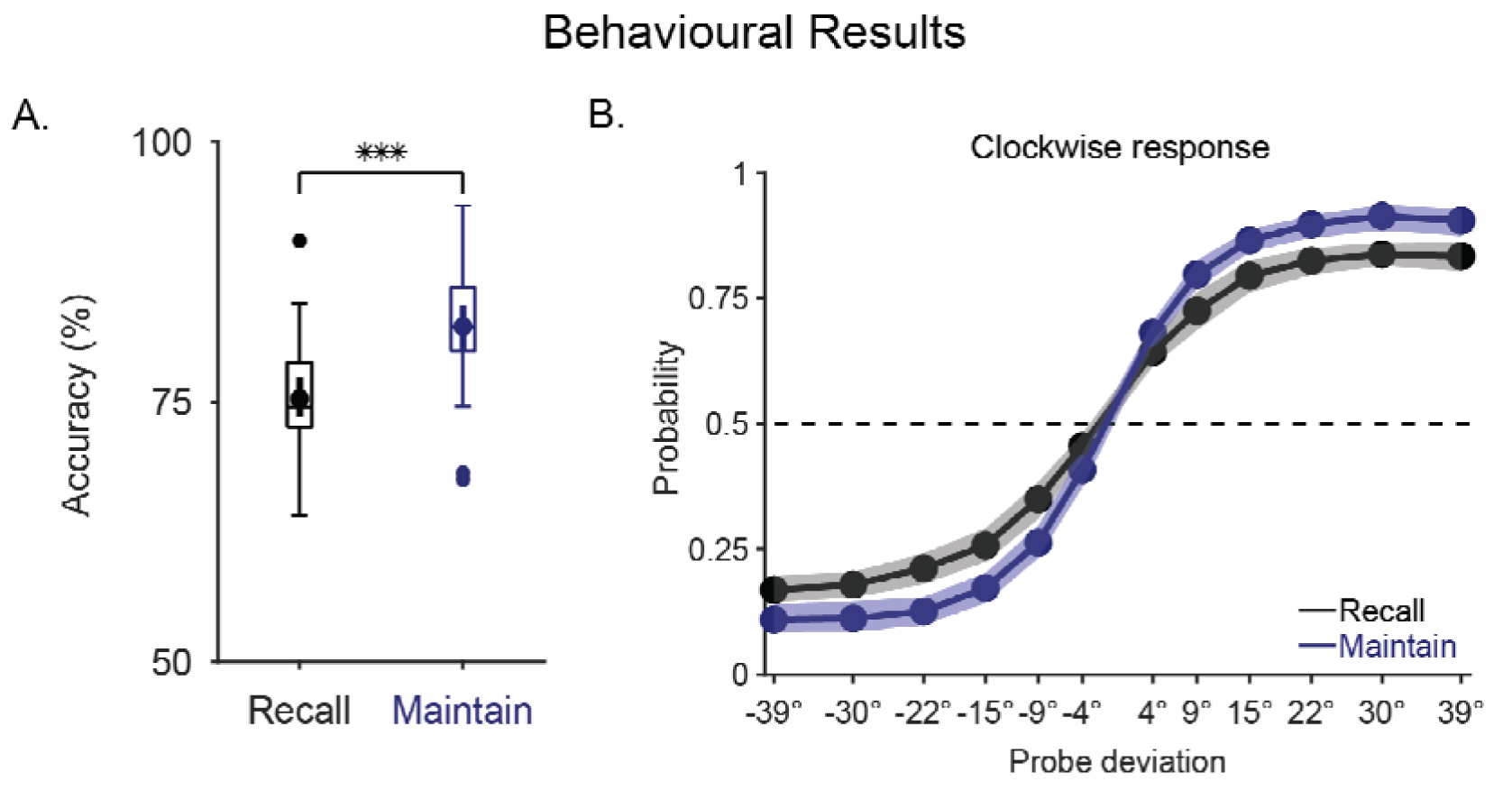
Behavioral accuracy (A) and Probability of clockwise response (B) *Note.* A. Response accuracy for Recall (black) and Maintain (blue) conditions. The middle lines in the boxes mark medians. The borders of the boxes cover the 25^th^ and 75^th^ percentiles, and the whiskers indicate the 1.5 interquartile range. The dots in the middle of the boxplots mark the mean accuracy for each condition, and their whiskers show the 95% confidence interval of the mean. Asterisks indicate statistically significant differences between conditions (***, *p* < 0.001). B. Probability of “clockwise” responses as a function of probe deviation, relative to the memory item. The dots indicate the mean probability for clockwise responses at each item-probe deviation range. The shaded zone marks the 95% CI.

### Decoding Orientations

The initial test was whether the visual orientations could be decoded both after the onset of the visual memory items and after the presentation of their auditory pairs. Visually presented orientations (Maintain condition) were almost immediately decodable (Fig. 3A blue line, 46 – 1694 ms relative to stimulus onset, *p* < 0.001, *two-tailed, cluster corrected*). Interestingly, we observed successful decoding after the auditory stimuli (Recall condition) early on as well (Fig. 3A black line, 46 – 1700 ms relative to stimulus onset, *p* < 0.001, *two-tailed, cluster corrected*). Crucially, a parametric relationship was apparent from onset in the Maintain condition (Fig. 3B, bottom), reflected by a gradual change in similarity as a function of angular difference, whereas such a parametric relationship was not present until later in the Recall trials (Fig. 3B, top). This suggested that the orientation memories might have been activated over time, even though the decoding algorithm could classify trials successfully based on the auditory cues that the orientations were paired with. This was also confirmed by the visualization of the representational code in state space, using EEG data from 100 to 400 ms post-stimulus (Fig. 3C). In Maintain trials the orientation stimuli could be clearly distinguished on two dimensions, whereas the same dimensions did not discriminate orientations in Recall trials. This pattern reflected that overall distances between tones were equalized over orientations across all participants, whereas by design, for orientation stimuli the underlying parametric differences remained constant for all.

**Figure 3.**
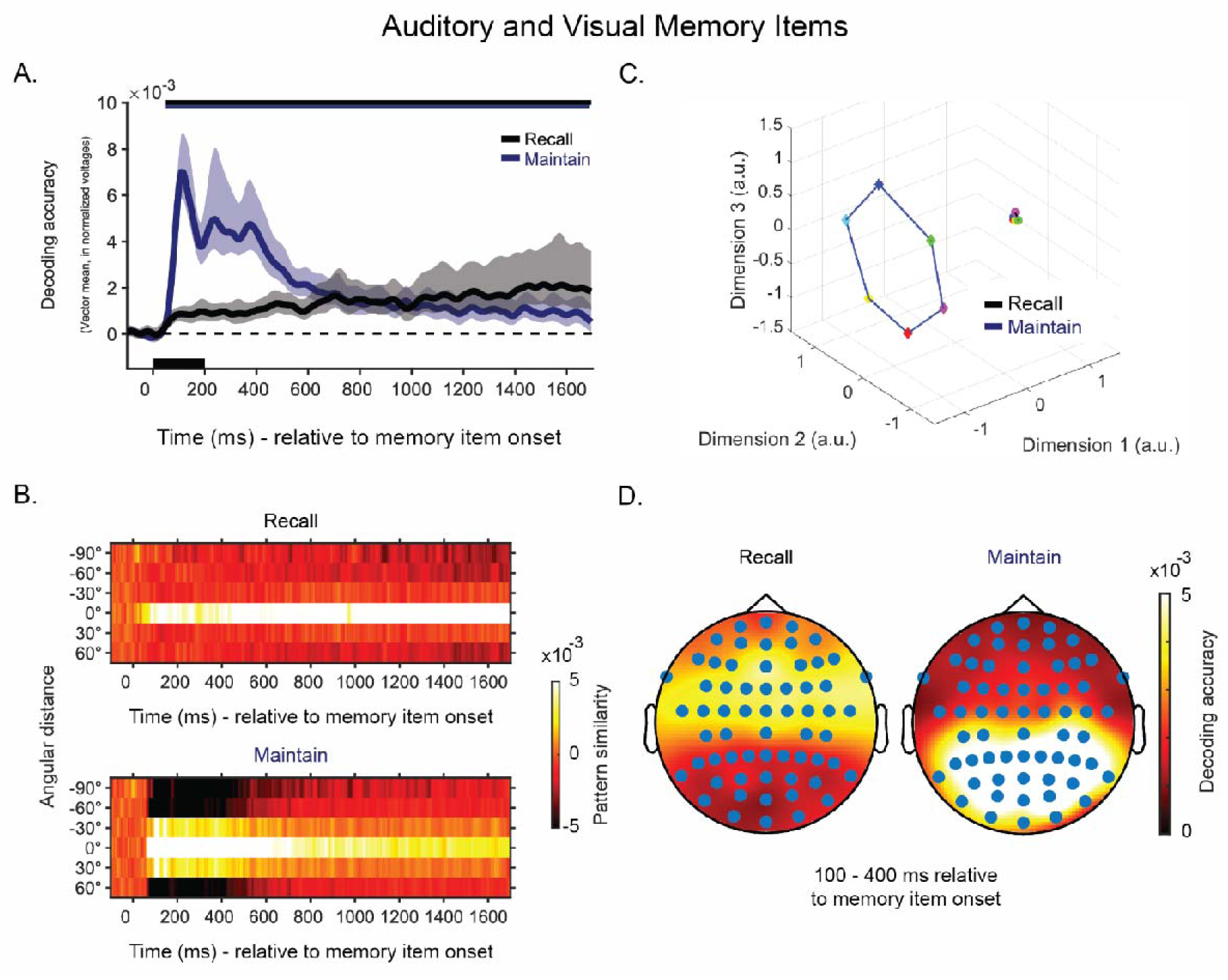
Orientation decoding after memory item presentation (A), Pattern similarities (B), State space for orientations (C) and Topographical distribution of representations (D). *Note.* A. Mean decoding accuracy for orientations in the Recall (black) and Maintain (blue) conditions as a function of time. The shaded zone around the line marks the 95 % C.I.. Horizontal lines at the top mark statistically significant time periods (p < 0.05, two-tailed, cluster corrected). B. Average similarity patterns (mean-centered, sign-reversed Mahalanobis distance) as a function of angular distance and time. C. Orientation code visualized in three-dimensional state space in the Maintain and Recall conditions. D. Topographical distribution of electrodes (with 2 neighbors) with statistically significant (p < 0.05) contributions to the decoding of the orientations in Recall and Maintain trials. Lighter colors reflect higher decoding accuracy.

Topographical analyses revealed that the trial-specific orientation memories could be inferred from all electrodes, but the main contribution was from the posterior electrodes in the Maintain condition (Fig. 3D, right), whereas central electrodes, nearer to the auditory cortices contributed most in Recall trials (Fig. 3D, right). The decoding of orientations thus seemed to rely on the actual, visual orientation representation in the Maintain trials, whereas in Recall trials it was the auditory stimulus associated with the orientation that was decoded, at least during the relatively early 100 – 400 ms interval presently considered.

The next analyses investigated if the auditory impulse allowed successful decoding of the orientation memories. Already at Impulse 1, the auditory impulse revealed orientation memories for a brief period in the Recall condition (Fig. 4A, black line, 116 – 180 ms relative to impulse onset, *p* = 0.046, *two-tailed, cluster corrected*). Conversely, the auditory impulse did not evoke statistically significant decoding in Maintain trials. In line with this, pattern similarity observed for target items was high in Recall trials, but not in Maintain trials (Fig. 4B). Topographical plots showed that even though time-course decoding was successful using the posterior electrode set, frontal areas actually reflected stronger representations for trial-specific memories in Recall trials (Fig. 4C).

**Figure 4.**
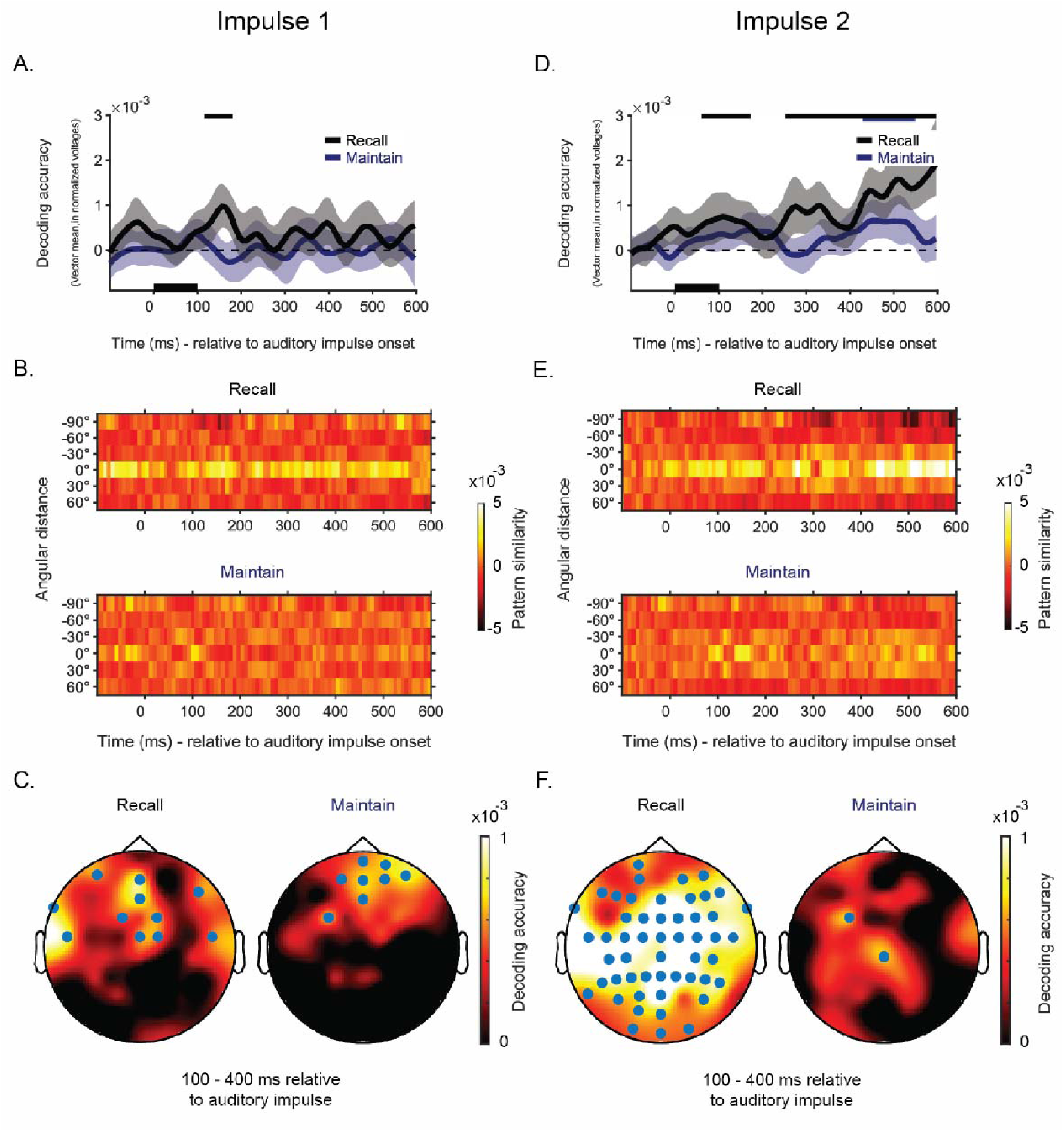
Time-course decoding at auditory Impulse 1 (A) and 2 (D), Similarity patterns (B and E), Topographic plots of auditory Impulse 1 (C) and 2 (F). *Note.* A & D. Mean decoding accuracy for orientations in Recall (black) and Maintain (dark blue) trials as a function of time, at the first (A) and second auditory impulse (D). The shaded zone around the line marks the 95 % C.I.. Horizontal lines at the top marks statistically significant time periods (p < 0.05, two-tailed, cluster corrected). B & E. Average similarity patterns (mean-centered, sign-reversed Mahalanobis distance) as a function of angular distance and time at the first (B) and second impulse (E). C & F. Topographical distribution of electrodes (with 2 neighbors) with statistically significant (p < 0.05, one-tailed) contributions to the decoding of the orientations in Recall and Maintain trials at Impulse 1 (C) and 2 (F). Lighter colors reflect higher decoding accuracy.

At Impulse 2, there was stronger and more consistent decoding after the auditory impulse in Recall trials (Fig. 4D, black line, 68 – 172 ms relative to impulse onset, *p* = 0.039, and 252 – 600 ms relative to impulse onset, *p* < 0.001, *two-tailed, cluster corrected*). Decoding in Maintain trials was also significant (Fig. 4D, blue line, 428 – 548 ms relative to impulse onset, *p* < 0.001, *two-tailed, cluster corrected*), although this fell outside the critical window, where the impulse effect is normally observed (Wolf et al., 2015; 2017; 2020a). Pattern similarity was highest for the target item and appeared parametric during periods with high decoding (Fig 4E). Topographical plots covering the critical time window showed that in Recall trials, memory-related activity could be decoded in a large area, including the central areas associated with audition as well as the posterior regions associated with vision (Fig. 4F, left). Conversely, in the Maintain condition, only two central electrodes showed significant decoding.

These results show that the learned associations between tones and orientations enabled auditory impulses to read out the visual working memory network. Successful decoding of visual memories was only possible when these memories were retrieved following the presentation of their learned auditory counterparts, whereas apart from a relatively late and weak effect outside of the typical critical post-impulse period, purely visually-driven memories did not respond to cross-modal pinging.

We also assessed orientation decoding from visual impulse-driven activity. At Impulse 1, both in the Maintain condition (Fig. 5A, blue, 28 – 380 ms relative to impulse onset, *p* < 0.001, *two-tailed, cluster corrected*), and in the Recall trials (108 – 524 ms relative to impulse onset, *p* < 0.001, *two-tailed, cluster corrected*), decoding was successful. At Impulse 2, the visual impulse led to even higher decoding in both the Maintain (Fig. 5D, blue, 148 – 192 ms relative to impulse onset, *p* < 0.001, *two-tailed, cluster corrected*) and in the Recall condition (Fig. 5D, black, 92 – 600 ms relative to impulse onset, *p* < 0.001, *two-tailed, cluster corrected*). Pattern similarity matrices for both conditions are presented in Figure 5B and 5E. These matrices show that especially when the decoding accuracy peaked, a parametric-looking relationship was observed for orientation memories. Topographical plots (Fig. 5C and 5F) revealed that in all conditions the visual impulse evoked a response in predominantly posterior regions, so that electrodes above these regions contributed highly to the decoding of orientation memories.

**Figure 5.**
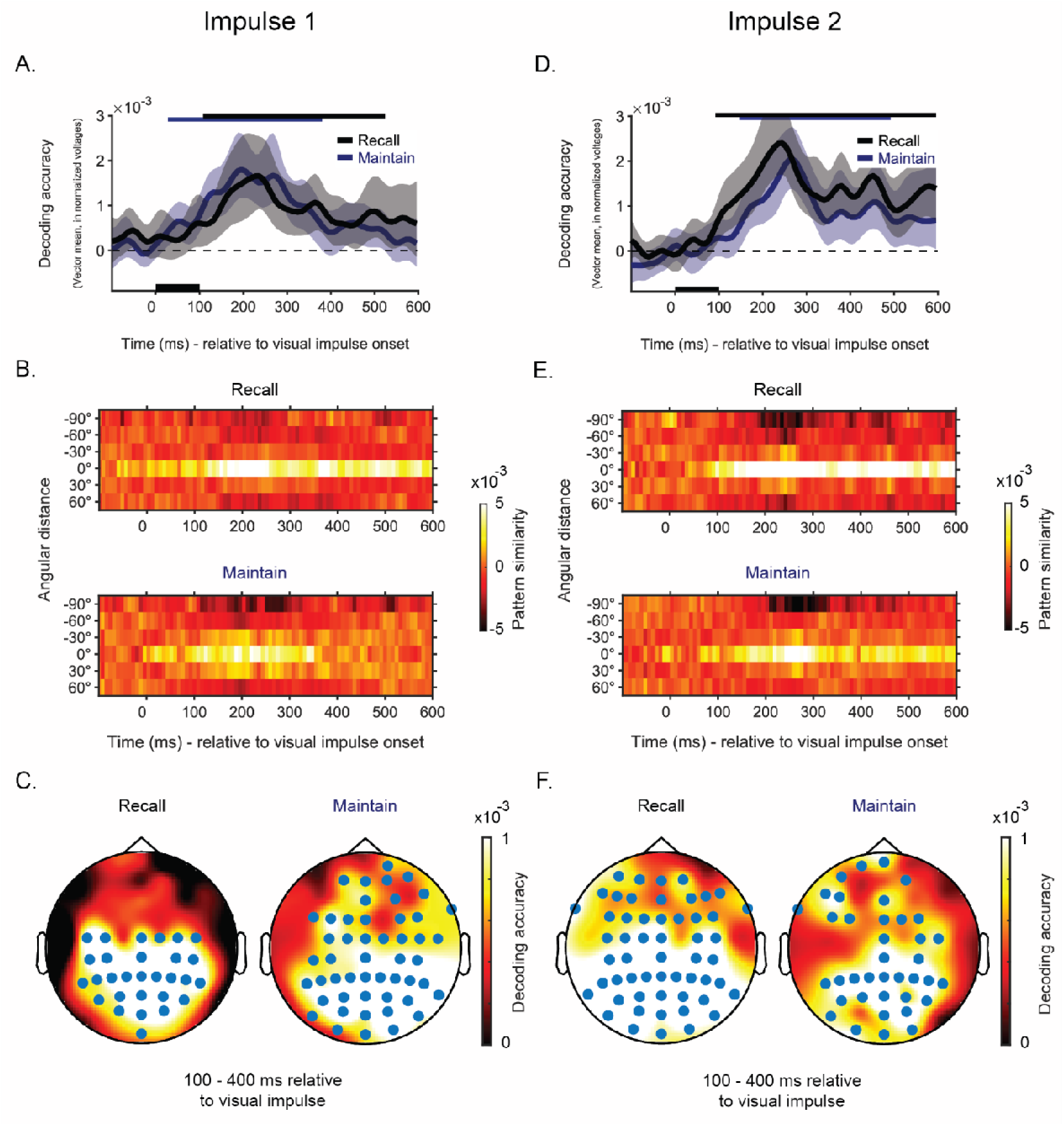
Time-course decoding at visual Impulse 1 (A) and 2 (D), Similarity patterns (B and E), Topographical plots of auditory Impulse 1 (C) and 2 (F). *Note.* A & D. Mean decoding accuracy for orientations in Recall (black) and Maintain (dark blue) trials as a function of time, at the first (A) and second auditory impulse (D). The shaded zone around the line marks the 95 % C.I.. Horizontal lines at the top marks statistically significant time periods (p < 0.05, two-tailed, cluster corrected). B & E. Average similarity patterns (mean-centered, sign-reversed Mahalanobis distance) as a function of angular distance and time at the first (B) and second impulse (E). C & F. Topographical distribution of electrodes (with 2 neighbors) with statistically significant (p < 0.05, one-tailed) contributions to the decoding of the orientations in Recall and Maintain trials at Impulse 1 (C) and 2 (F). Lighter colors reflect higher decoding accuracy.

These results show that visual impulses are not only useful to reveal recently encoded visual working memory content during a delay (e.g., Wolff et al., 2020b), but that they can also reveal retrieved (activated) visual long-term memories.

We wanted to know whether the memory representations might differ for Recall and Maintain conditions. This was tested by assessing the generalization of the memory representations across conditions, collapsed over same-modality impulses. For the visual impulse, memory representations for orientations proved to be similar across Recall and Maintain conditions (Fig. 6A, pink, non-parametric permutation test (*n_perm_* = 100 000), *p* < 0.001, *two-tailed*). Conversely, orientation decoding did not generalize across conditions for the auditory impulse (Fig. 6A, light blue, non-parametric permutation test (*n_perm_* = 100 000), *p* = 0.297, *two-tailed*). The similarity patterns revealed that the parametric-looking relationship between memory items persisted in both conditions following the visual impulse. This was not the case when the conditions were compared after the auditory impulse, reflecting that the auditory impulse likely did not perturb the same network in both conditions (Fig. 6B).

**Figure 6.**
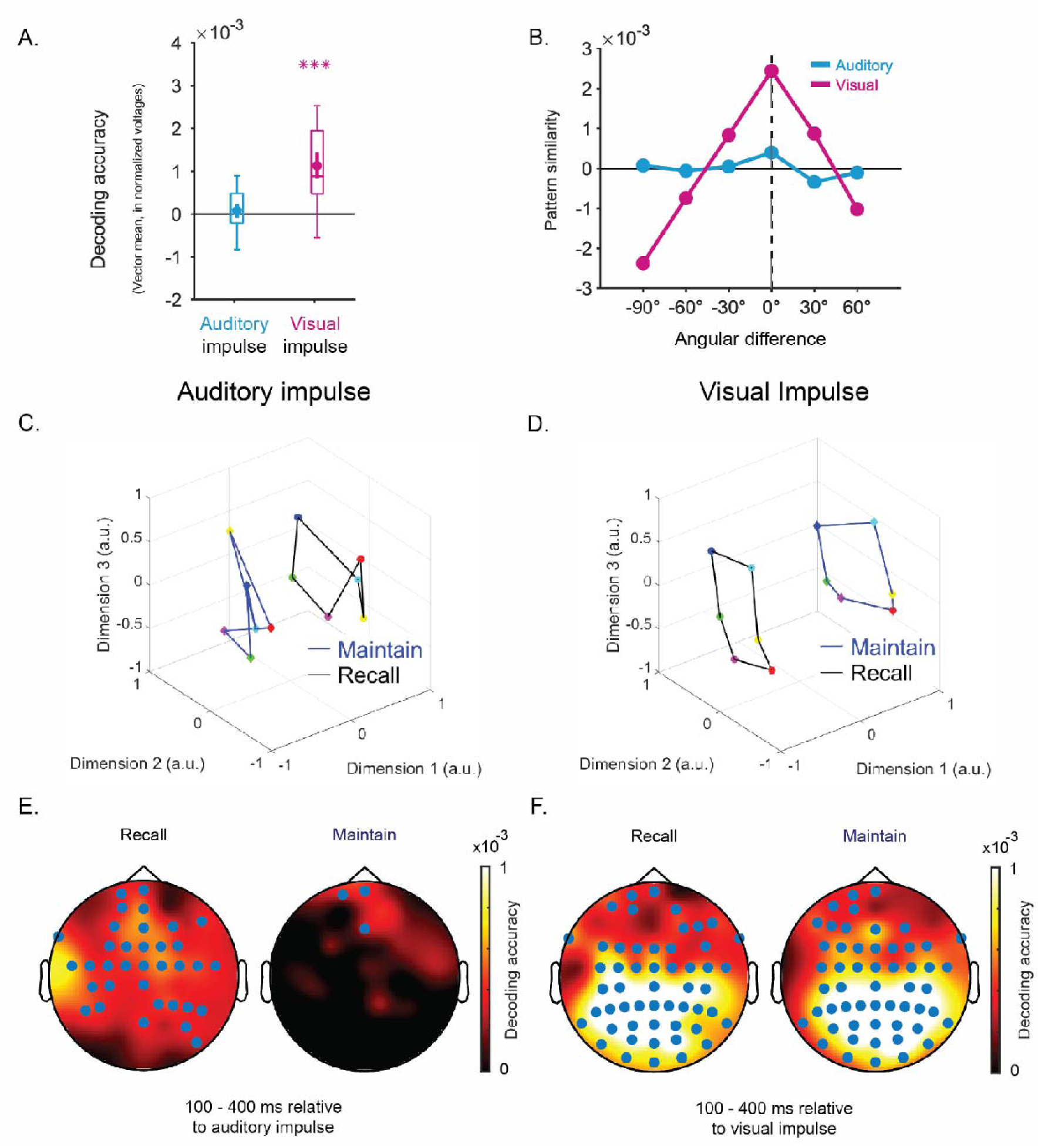
Generalization of memory representations across conditions (A) and associated pattern similarity (B), Representation code in state space after auditory and visual impulse presentation (C and D), Topographical plots for auditory (E) and visual impulses (F). *Note.* A. Boxplot showing cross-generalization of orientation coding after auditory (light blue) and visual impulses (pink), by training and testing independently across Recall and Maintain conditions, using the 100-400 ms critical window collapsed over the first and second impulses. The middle lines in the box mark the median. The borders of the boxes cover the 25^th^ and 75^th^ percentiles, and the whiskers indicate the 1.5 interquartile range. The dot in the middle of the boxplot marks the mean decoding accuracy for each condition, and its whiskers show the 95% C.I.. Asterisks indicate statistically significant decoding (***, p < 0.001, two-tailed). B. Similarity patterns of the neural code across different experimental conditions, after the auditory (blue) and visual (pink) impulse. C-D. Orientation code visualized in 3-dimensional state space for Maintain and Recall conditions, following the auditory (C) and visual impulse (D). E-F. Topographical distribution of electrodes (with 2 neighbors) with statistically significant (p < 0.05, one-tailed) contributions to the decoding of the orientations in the Recall and Maintain conditions, following the auditory (E) and visual impulse (F). Lighter colors indicate higher decoding accuracy.

Visualizing these representational codes in state space (100-400 ms post-impulse), yielded a circular relationship across orientations after the auditory impulse, but only in the Recall condition (Fig. 6C). The emergence of this pattern, coupled with the complete absence of it after the presentation of the auditory item (cf. Fig. 3C), provided strong evidence that the neural pattern at this stage reflected the retrieved orientations, rather than anything related to the auditory stimulus itself. Further, a very similar relationship was clearly present in both Recall and Maintain conditions after the visual impulse (Fig. 6D). Finally, searchlight analyses suggested that the auditory impulse effect (in the Recall trials) seemed to include a fairly distributed set of electrodes (Fig 6E.). Conversely, the visual impulse effect seemed heavily driven by occipital brain regions, regardless of whether there was recall of an orientation, or just maintenance (Fig. 6F), thus suggesting that the two impulse modalities might use different access pathways.

The temporal stability of the neural signal was assessed by testing the neural pattern at each time point over all time points within the epoch. In Recall trials, the neural pattern was initially dynamic. This dynamic state was reflected by the higher decoding observed when training and testing on the same time point relative to other possible time points (Fig. 7A). This dynamic pattern, which was likely associated with the encoding of the auditory item, was overtaken by a pronounced static pattern starting from 600 ms, possibly reflecting the retrieval of orientation memories. This static pattern persisted over the remainder of the period. In Maintain trials, the pattern also had a dynamic property in the early period of the epoch (Fig. 7B). However, there was considerable evidence for stable coding as well, which seemed to start already from 400 ms post-stimulus.

**Figure 7.**
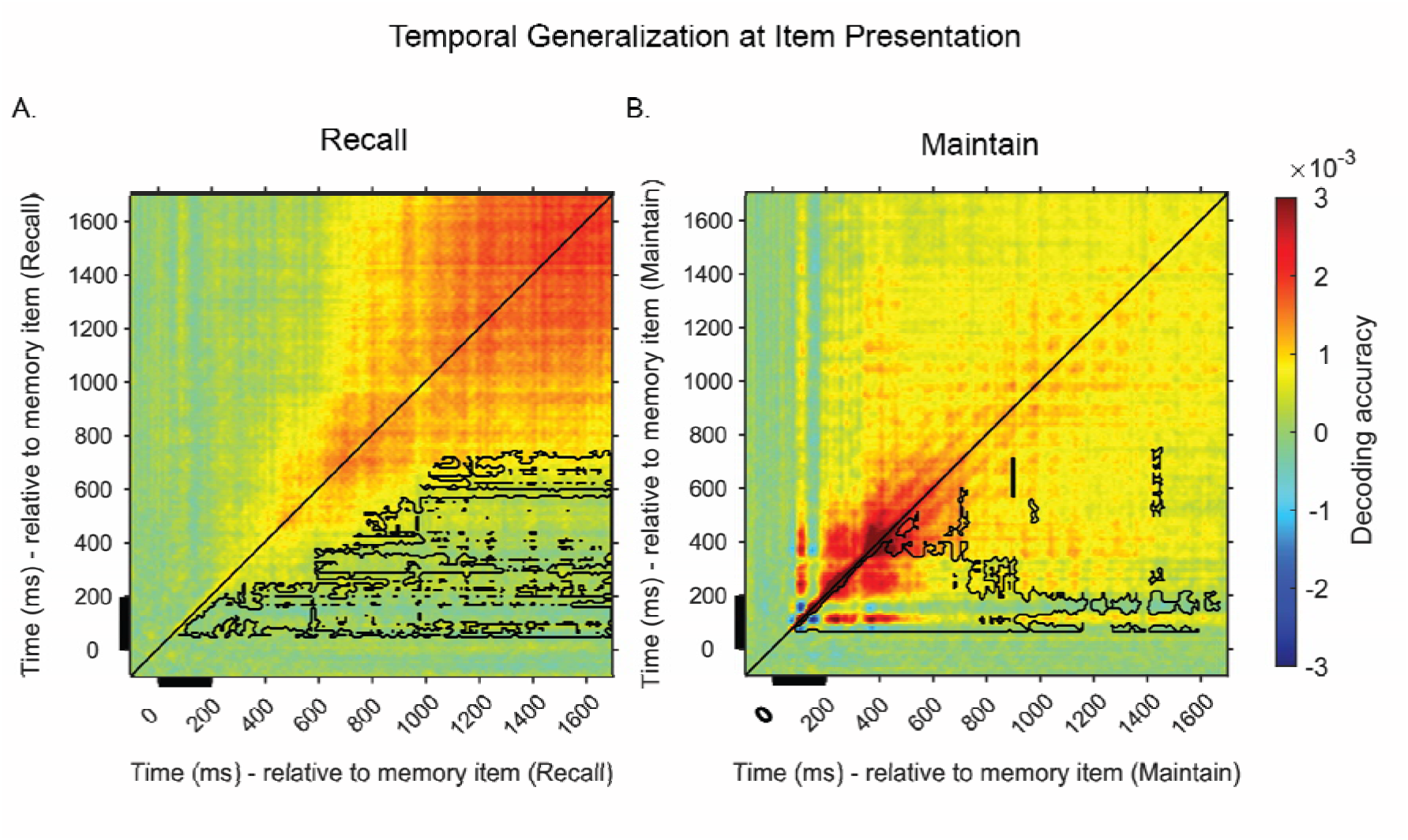
Cross-temporal generalization of memory representations across conditions (A) Recall condition (B), Maintain condition. *Note.* Cross-temporal decoding matrices during and after item presentation. A. Cross-temporal generalization in Recall trials B. Cross-temporal generalization in Maintain trials. Decoding accuracy is represented by color intensity. Black lines mark the edges of the time-zone during which there was statistically significant lower decoding relative to the diagonal (same train and test time points; *p* < 0.05, *cluster corrected*). Black rectangles mark the presentation of the memory items.

The temporal dynamics of the neural representations following the impulse presentation also reflected co-existing dynamic and static patterns. Following the presentation of the visual impulse in the Recall condition (Fig. 8A, left), the decoding accuracy along the diagonal was significantly higher than at other time points, especially in the critical time-window of 100 - 400 ms, although off-diagonal decoding accuracy was also clearly present across the epoch. Similarly, in the Maintain condition, the visual impulse revealed higher decoding on the diagonal, indicating a dynamic code, but off-diagonal decoding was also quite high (Fig. 8A, middle). When training on Recall and testing on Maintain trials, the generalization across conditions seemed to rely most on a static pattern (Fig. 8A, right). For the cross-modal auditory impulse, decoding also cross-generalized over time, reflecting a stable representational code (Fig. 8B). These outcomes suggest that cross-modal retrieval in WM, as reflected by the auditory impulse response, relies on a predominantly stable code, whereas the visual impulse response showed more evidence for (initial) dynamic coding.

**Figure 8.**
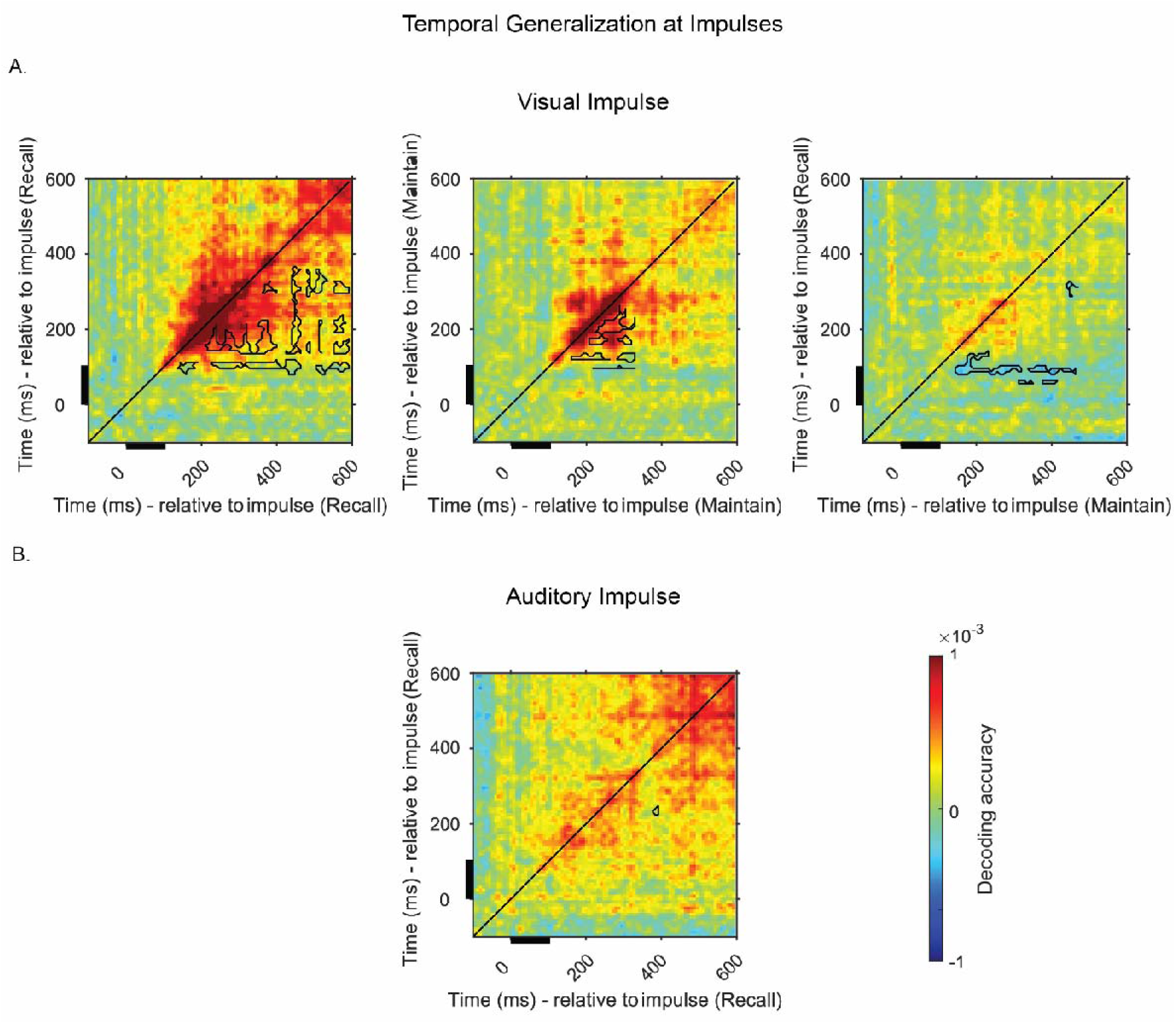
Cross-temporal generalization of memory representations at (A) visual impulse and (B), auditory impulse for Recall and Maintain conditions. *Note.* Cross-temporal decoding matrices during and after impulse presentation. A. Decoding after the visual impulse (collapsed over Impulse 1 and 2) by training and testing on all time-point combinations in Recall (left) and Maintain (middle) conditions, and cross-temporal generalization over Recall and Maintain conditions (right). B. Cross-temporal generalization after the auditory impulse in the Recall condition. Warm colors reflect higher decoding accuracy. Black lines mark the edges of the time-period in which equivalent vertical or horizontal timepoints contain statistically significant lower decoding, relative to the decoding at the same time point (on the diagonal) (p < 0.05, cluster corrected). The black rectangles mark the presentation of the impulse.

## Discussion

The principal aim of the current study was to test whether learned information in long-term memory can provide cross-modal access to working memory. To this end, we attempted to decode a single visual working memory item from the neural activity evoked by an auditory impulse stimulus. Previous research has shown that without learning, an auditory impulse can only reveal auditory memories, not visual ones (Wolff et al., 2020b). We found that by learning to pair visual orientations with auditory tones, these orientations could be decoded from auditory impulses presented during working memory maintenance.

The cross-modal perturbation of visual orientations in working memory only occurred when retrieval of these items was required through their paired auditory counterparts. On trials in which the orientation stimuli were shown directly, the orientations could not be decoded from the auditory impulse within the critical period during WM maintenance. The mere passive presence of the auditory-visual pairs in long-term memory was thus by itself not sufficient to bridge the different sensory modalities; active retrieval of the relevant information was needed. Finally, we observed that cross-modal access to WM tended to produce more stable representational coding during encoding and maintenance. Our findings have implications for (neural) models of working and long-term memory, as well as for the impulse perturbation technique, which are discussed below.

### Brain areas involved in cross-modal WM

Learning of cross-modal stimuli reportedly follows the co-activation of unimodal sensory regions, together with multimodal regions like the occipitotemporal junction and parahippocampal gyrus, which realize the binding of modalities (Calvert, 2001; Tanabe et al., 2005). In addition to the indirect links involved in the integration and association of auditory and visual stimuli, direct neural links from auditory cortex to the visual cortex likely exist as well (Garner & Keller, 2021). Earlier PET and fMRI work that consistently paired an auditory tone with a visual stimulus has also shown cross-modal activation of the visual cortex by auditory stimulation (McIntosh et al., 1998), and the reverse when visually displayed musical notes were used for auditory imagery and pairs of objects and sounds (Hoppe et al., 2014; Suchan et al., 2006). The present results are in principle compatible with the idea that sensory regions played an important role in our cross-modal WM task. The topographical searchlight analyses did not provide evidence that the effects should be localized very differently either.

Nevertheless, apart from sensory regions, the involvement of the hippocampus in the current task may be expected, considering its role in learning (Morgado-Bernal, 2011), declarative (Eichenbaum, 2001) and spatial memory (Stella et al., 2011). Because the auditory impulse was only effective during retrieval, it seems unlikely that it reflected hippocampus-mediated long-term codes, as these would have been present during Maintain trials also. Therefore, we believe the representations in the current task probably originate more directly from links between auditory and visual cortices, with possible mediation of multimodal areas.

### Impulse perturbation and short-term synaptic plasticity

Regardless of the specific brain areas that are implicated, representations in WM could be formed not only by persistent neuronal firing, but also by short-term synaptic plasticity (Barak & Tsodyks, 2014; Hempel et al., 2000; Mongillo et al., 2008; Stokes, 2015). The synaptic view holds that the encoding of a stimulus temporarily changes the synaptic connections in the memory network (e.g., through modulations of presynaptic calcium ions), thereby producing a transient and activity-quiescent memory state. This state can be perturbed and thereby read out by driving generic activity through the network, as is achieved by the presentation of an impulse stimulus (Wolff et al., 2017; 2020b). The current experiment indeed replicated this impulse effect. At the same time, considering that long-term memories are likely formed through synaptic alterations (Abraham, Jones, & Glanzman, 2019; Bear, 1996; Hebb, 1949; Stent, 1973), there was little evidence that such more durable synaptic changes were sensitive to impulse perturbation. Although one might doubt whether impulse perturbation could reveal long-term memories in general, an impulse response might be conceivable when the long-term memories are currently task-relevant, as they were in the present study. However, the lack of a compelling auditory impulse effect in the Maintain condition that we presently observed would seem to contradict this idea. Our data cannot arbitrate as to why this might be the case. It is possible that long-term memory is simply further from the surface of mind and brain (e.g., hippocampal) than WM is. In a similar vein, a minimal level of activity may be needed for a robust impulse effect, which might be afforded by periodic refreshing in WM (e.g., in the alpha band; Barbosa et al., 2021). This question must be left for future research.

### WM as activated LTM

In functional terms the current outcomes provided support for models that posit that WM might be conceived of as an activated portion of long-term memory. We observed that the visual impulse revealed the memoranda both when orientations were visually encoded, as well as when an auditory cue guided retrieval, replicating earlier robust visual impulse effects (Wolff et al., 2017, 2020a; 2020b). Importantly, the neural code for orientation memories generalized across both conditions, revealing that visually encoded memories relied on similar representations as the memories that were retrieved from long-term memory. This finding supports state-dependent memory models, such as the embedded-processes model (Cowan, 1999; Adams et al, 2018). This model contains a unitary memory system, in which WM corresponds to a portion of the memory network that is activated by the focus of attention, and in which no consequent change in representational code between dormant and active states would be expected.

The sensory-recruitment hypothesis extension of the embedded-processes model further proposes that the neural networks and coding scheme associated with the perception of a stimulus is also employed during working memory maintenance (Awh & Jonides, 2001; D’Esposito & Postle, 2015; Scimeca et al., 2018). According to these views, the representation of memoranda is reflected by a static stable neural code under identical selection and activation levels over short and long temporal durations. In the present study, we observed co-existing dynamic and static neural codes for orientation memories independent of their source (perceptual or from long-term memory). Although visual impulses in particular evoked an initial dynamic response, the generalization across conditions relied on the stable neural code. These results are thus largely in line with the embedded-processes model and the sensory-recruitment hypothesis, since both the visually encoded memories and those retrieved from long-term memory relied on similar, stable neural representations. Since the impulse effect is due to generic sensory stimulation, it also seems likely that these stable codes originated from sensory (visual) cortex.

## Conclusion

The current study showed that cross-modal functional connectivity underlying WM is subject to change by learning. These changes were large enough to allow auditory impulse perturbation of the visual working memory network, which was not observable before (Wolff et al., 2020b), supporting the utility of the impulse perturbation technique in investigating (cross-modal) WM. Active retrieval of the memoranda from WM seemed to be an important factor in our study, as the auditory impulse effect was not convincingly present when visual items were merely maintained after presentation, suggesting that the long-term links themselves, when passive, do not meaningfully contribute to this effect. Finally, we observed (partially) stable neural signatures independent of how the memoranda were accessed, suggesting commonality in the coding schemes for working and long-term memory representations.

## Acknowledgements

The authors would like to thank Kristin Langohr and Ania Stró_ż_yk for their help with data collection, and Mark Span for his help with the timing of the auditory stimuli.

## Supplementary Materials

Participants learned to pair the auditory and visual stimuli offsite, at a time and location of their own choosing, and using their own devices to run the program. There was no control over the sound levels of the auditory stimuli, screen resolution or visual stimulus size. Each participant ran a predesigned individual version of the OpenSesame experiment, so that tone-orientation pairings were evenly distributed across the participants. The program comprised two parts. The first part was a study session, and consisted of a flashcard procedure with paired presentation of auditory and visual stimuli from low to high auditory frequencies. The second part was a practice session, in which randomly chosen auditory stimuli prompted the recall of their visual pairs.

In the study part, a trial consisted of the presentation of the entire stimulus set, formed by 6 tones and 6 orientations. A trial began after participants pressed the Spacebar. A warning message (“Get Ready”) was presented in Serif size 32 font, at the center of the screen for 200 ms, followed by the presentation of a fixation dot (white edges surrounding a black dot) for 200 ms. Next the first pure tone and its associated orientation grating were presented simultaneously for 200 ms. The remaining tone-orientation pairs were consecutively presented, separated by a blank interval of 1000 ms. All pairs were presented following the order of the tones from low to high frequency. At the end of the presentation sequence participants could press the Spacebar to repeat the sequence as many times as they wished or press “C” to begin the practice part.

During practice, participants were asked to recall the orientations that were associated with the tones. A practice trial began by participants pressing “M” which was followed by the presentation of the warning signal. Following the presentation of the fixation dot for 200 ms, a randomly selected auditory tone was presented for 200 ms. This was followed by a 1000 ms delay, during which only the fixation dot was present on the screen. Then a sine-wave grating with a random orientation was displayed at the center of the screen. By moving the mouse horizontally and vertically around the center of the screen, the participants could rotate the grating to designate the correct orientation associated with the auditory stimulus. The response was submitted by a click of the left mouse button. Following the response, a feedback screen was presented. The feedback consisted of a white line depicting the orientation of the response given, and a red line depicting the correct orientation. The mean deviation between response and actual orientation was calculated throughout the practice trials and shown to the participants at the end of a block of 12 practice trials.

The frequency and the duration of the study and the practice sessions completed by the participants, until they entered the screening procedure, was self-determined. Participants were informed that an average of 13° deviation in reproduction of the orientation over a period of 12 trials would likely reflect adequate learning of the stimuli, which was a value observed in a previous pilot experiment. It was possible for participants to take the screening test and then re-study in order to take part in the EEG session.

